# Insights into Hepatitis C Transmission in Young Persons who Inject Drugs: Results From a Dynamic Modeling Approach Informed by State-Level Public Health Surveillance Data

**DOI:** 10.1101/193185

**Authors:** Rachel E Gicquelais, Betsy Foxman, Joseph Coyle, Marisa C Eisenberg

## Abstract

Rising use of heroin and prescription opioids are major contributors to increases in Hepatitis C Virus (HCV) incidence in US young adults since the late 1990s. How best to interrupt transmission and decrease HCV prevalence in young persons who inject drugs (PWID) is uncertain, but modeling studies in older populations support interventions that increase HCV treatment among all PWID. We developed a transmission model of young (aged 15-30 years) PWID, which we fit to state-level US HCV surveillance data, and simulated the potential impact of primary (reducing injection initiation), secondary (increasing cessation, reducing injection partners, or reducing injection drug use relapse), and tertiary (HCV treatment) interventions on incident and prevalent HCV cases. Interventions with primary prevention initiatives (reducing injection initiation) yielded concurrent reductions to HCV incidence and prevalence. Treatment of former PWID led to prevalence reductions but did not reduce incidence. Treatment of current and former PWID without other interventions led to incidence reductions in scenarios with high injection initiation rates, high syringe sharing, and low relapse rates after injection cessation. While these results are specific to Michigan, our approach could be applied in other states conducting HCV surveillance to identify local-level intervention opportunities.

## INTRODUCTION

The epidemiology of hepatitis C virus (HCV) in the United States (US) has changed dramatically over the last decade. Increases in HCV incidence among young persons aged approximately 15-30 years have been linked to increases in opioid and injection drug use (IDU), which have been especially large in rural communities (1–5). In the US, up to 2.6% of adults have injected drugs in their lifetime and 38-48% of US persons who inject drugs (PWID) have HCV infection, although the prevalence among young PWID is not generally well-characterized (6–10). After decades of chronic infection, HCV leads to liver-associated morbidity and mortality (e.g. cirrhosis and hepatocellular carcinoma) (11,12).

In the US, transmission modeling studies have shaped HCV screening, treatment, and prevention policies by increasing understanding of HCV transmission dynamics, forecasting prevalence of HCV-related liver diseases, and simulating the impact, costs, and benefits of highly effective directly acting antiviral (DAA) treatments among PWID and other groups disparately burdened by HCV (11,13–18). Several different modeling approaches support the cost-effectiveness of treating PWID with DAAs to interrupt HCV transmission, a strategy known as “treatment as prevention” (14,15,17,19–42). One of the few modeling studies focused on young US PWID (<30 years) showed that, in Chicago, a treatment rate of just 5 per 1,000 young PWID could halve HCV prevalence over 10 years in this age group (from 10% to 5%) (14). While available for Chicago, estimates of HCV prevalence specific to young PWID are not routinely available at the state or local-level throughout the US, limiting the ability to forecast prevalence and evaluate the impact of ongoing interventions. We demonstrate here that routine HCV public health surveillance data collected as part of nationally notifiable and state-reportable condition surveillance might be used for this purpose.

HCV incidence and prevalence increases in young adults and adolescents were first identified using HCV public health surveillance data (2,43,44). As part of HCV surveillance, laboratories and physicians report positive HCV lab results to state health departments, who apply standard case definitions to stage HCV as acute or chronic (45,46). Underreporting of HCV cases limits use of surveillance data for purposes other than description and outbreak monitoring (2). Klevens et al. recently estimated the magnitude of acute HCV under-reporting at approximately 12.3-16.8 cases per case reported (47). In addition to under-reporting, there is high variability in capacity to collect risk factor or demographic information, trace contacts, and connect persons with HCV infection to testing and treatment (2,4). These limitations have discouraged use of public health surveillance data for HCV transmission modeling.

We developed an HCV transmission model of young PWID in Michigan fit to HCV surveillance data for the state of Michigan (48) that adjusts for case under-reporting and incorporates parameter uncertainty through Latin hypercube sampling of the parameters, a form of stratified random sampling. Model simulations evaluate interventions in a counterfactual framework, including primary prevention (reduced injection initiation), secondary prevention (behavioral initiatives), and tertiary initiatives (HCV treatment), for reducing HCV prevalence and incidence. An evaluation of model fit to data given parameters in the literature is used to identify the consistency of parameter ranges with surveillance data and identify scenarios under which certain interventions may be optimal. This modeling framework could be applied to HCV surveillance data from other states and/or adapted for use in other nationally or state-notifiable conditions.

## METHODS

A detailed discussion of the model structure, parameters, initial conditions, surveillance data, and parameter estimation process is available in the Web Methods supplement. Matlab code for model simulation and R (R Foundation for Statistical Computing, Vienna, Austria) code for figures is freely available at https://github.com/epimath/Hepatitis-C-in-Young-PWID.

### Model Structure

An HCV ordinary differential equation (ODE) transmission model of PWID with preferential age mixing was developed in Matlab R2016a (The MathWorks Inc, Natick, MA). The model consists of 9 states per age group, with age groups of 15-19 years, 20-24 years, and 25-30 years (Figure 1). Individuals age through groups and transition through a series of compartments within age classes: uninfected, non-PWID (Z_*i*_), uninfected current or former PWID (S_*i*_ or S_N*i*_, respectively), acutely infected current or former PWID (A_*i*_ or A_N*i*_, respectively), chronically infected current or former PWID (C_*i*_ or C_N*i*_, respectively), and treated current or former PWID (T_*i*_ or T_N*i*_, respectively).

**Figure 1.**
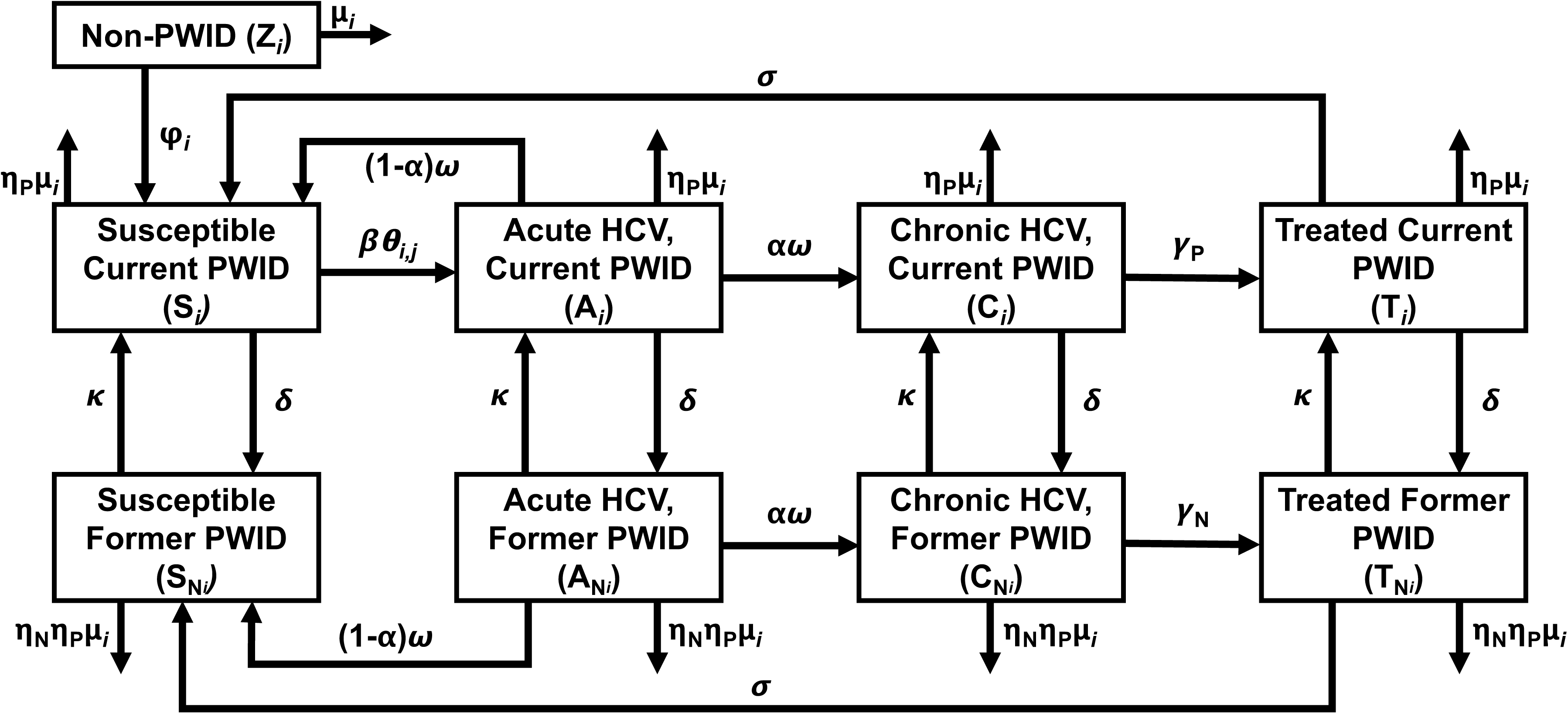
Compartmental Model of HCV Transmission among Young PWID in Michigan. States and parameters controlling flows between compartments of an HCV transmission model among young PWID in Michigan. Non-PWID (Z_*i*_) begin injecting drugs (S_*i*_) at a rate ϕ_*i*_ and acquire acute infection (A_*i*_) through effective contact with an HCV-infected PWID (A_*j*_ or C_*j*_) of the same or discordant age (**Θ**_*i,j*_) at a transmission rate (β). Chronic infection (C_*i*_) develops at a rate ω among a proportion of acute cases, α, and resolves in the remaining 1-γ acute cases. Chronic infection can be treated (T_*i*_) at a rate γ_P_, for a duration of σ^-1^ years, after which point susceptibility to reinfection ensues. PWID stop injecting drugs at a rate δ and transition to former PWID states (denoted by State_N*i*_). Former PWID can begin injecting drugs after a period of abstinence at a rate κ. Death occurs at a rate μ_*i*_, which is elevated among current PWID by a factor η_P_. Former PWID mortality rates are less disparate from non-PWID by multiplying the current PWID mortality increase factor by a protective factor η_N_. Subscript *i* denotes parameter or state age class (1=15-19 years, 2=20-24 years, 3=25-30 years) and subscript *j* denotes the age group of contacts from whom susceptible current PWID (S_*i*_) can acquire infection. Individuals move through age groups based on the duration predicted by the age range captured in each group (v_i_, not depicted for simplicity) and new 15 year-olds are added to the non-PWID compartment each year at a rate (v_0_Z_0_, not depicted for simplicity).

Individuals acquire HCV by injecting drugs; other transmission modes (e.g. perinatal acquisition, unregulated tattoos, sexual transmission) are not considered. Non-PWID initiate IDU at an estimated injection initiation rate, ϕ_*i*_, calibrated to fit HCV surveillance data from an initial value based on the system of equations at steady state (see Web Methods). Susceptible current PWID acquire new infections at an estimated rate β through effective contact (i.e. syringe sharing) with an acutely (A_*j*_) or chronically (C_*j*_) infected current PWID in any age class. Contact rates between individuals in each age class are parametrized by a contact matrix, θ, adapted from social contact patterns in eight European countries (49) (Web Methods). Individuals in treatment compartments (T_*i*_ and T_N*i*_) do not transmit HCV (i.e. are assumed not to be sharing syringes). Individuals from any of the current PWID classes can stop injecting drugs and move to their adjacent former PWID class at a cessation rate δ while former PWID can begin injecting again after a period of injection abstinence and enter the current PWID class at a relapse rate κ.

### Surveillance Data, Parameter Estimation, and Parameter Sampling

The Michigan Department of Health and Human Services (MDHHS) receives reports of HCV diagnoses from healthcare providers and laboratories and stages cases as acute or chronic using standardized national case definitions (45,46). We obtained the number of newly identified acute and chronic HCV cases per year during 2000-2013 and adjusted for under-reporting to surveillance sources using a correction factor developed by Klevens et al. (47). While this correction factor was developed to scale acute HCV cases, the extension to newly reported chronic HCV is justified in persons aged 15-30 years. These individuals likely acquired infection relatively recently and chronic HCV is under-diagnosed and under-reported among adolescents and young adults.

To incorporate parameter uncertainty, Latin hypercube sampling with 5,000 simulations was used to draw a stratified random sample of parameter sets across plausible ranges gathered through literature review (see Web Methods). To optimize model fit to data, we estimated four unknown parameters (the transmission rate [β] and three age-specific injection initiation rates [ϕ_*i*_]) in each simulation using weighted least squares assuming normally distributed measurement error with variances proportional to the data. Residual sum of squares values after parameter estimation were used to select the best-fitting 50% of parameter sets to data. To determine if a certain range appeared more consistent with data, we plotted histograms by quartile of fit to the data to visually examine if fit differed along uniformly sampled parameter ranges. Parameter estimation and simulations were run using fminsearch and the ODE15S solver in Matlab.

### Potential Interventions

For each of the 50% best-fitting parameter sets (2,500 simulations), the model was re-simulated after scaling one or more parameters, which provided counterfactual estimates of expected prevalence and new chronic cases (our measure of incidence) in the presence of interventions. We simulated primary prevention interventions (reduced injection initiation [(ϕ_*i*_]), secondary prevention interventions (decreased syringe sharing [θ], decreased IDU relapse [κ], and increased cessation [δ]), and tertiary prevention interventions (treatment of former [γ_N_] and current PWID [γ_P_]) (50). Although parameters cannot be singly classified into primary, secondary, and tertiary (e.g. current PWID treatment is tertiary [treatment] and secondary [reduces transmission]), these terms classify interventions by their most immediate roles. Parameters were scaled at 10%, 20%, and 40%. We first simulated the expected impact of each intervention alone. Next, we simulated the expected impact of combinations of interventions, specifically to compare the added benefit of secondary interventions on top of primary versus tertiary approaches. Intervention results are summarized by comparing the expected case counts for incident and prevalent chronic HCV cases at year 14 of simulation compared to no intervention.

Two intervention sensitivity analyses were conducted. First, case counts were examined after adding mortality reduction interventions (η_N_ and η_P_) of 40%, which simulate, for example, real-world initiatives to reduce overdose among current PWID, such as bystander overdose response training. Second, treatment duration was compared at 12 weeks versus 1 year, representing treatment durations of currently used DAAs to interferon-based treatments used before 2012, respectively (12).

## RESULTS

We simulated the potential impact of primary (reducing injection initiation), secondary (increasing cessation, reducing injection partners, or reducing injection drug use relapse), and tertiary (HCV treatment) interventions on incident and prevalent chronic HCV cases among young (aged 15-30 years) PWID. The model fit well to Michigan HCV surveillance data, including for the subset of 2,500 parameter sets used to simulate interventions (Web Figure 1). Better-fitting parameter sets that were used in intervention simulations tended towards the lower sampled bounds of current PWID prevalence, cessation, and syringe sharing contacts among 15-19 year olds and towards the higher sampled bounds of lifetime PWID prevalence and several other contact parameters (Web Figure 2). The reporting rate had a slight tendency toward the minimum rate, closer to the estimated rate of acute HCV detection among PWID cases in Klevens et al. (47). Other parameters fit consistently throughout the sampled range (Web Figure 3).

**Figure 2.**
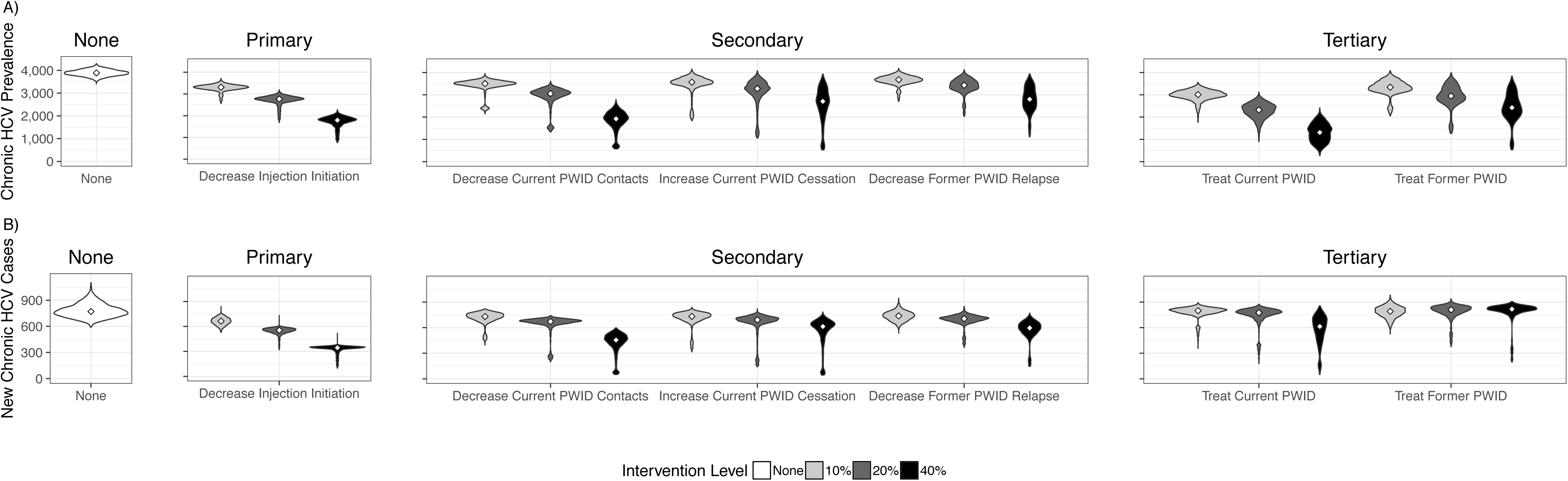
Counterfactual Simulation of Single Interventions. The distribution of predicted surveillance-detected HCV prevalence (top) and number of new chronic HCV cases (bottom) for each of 6 interventions among the best-fitting 50% of parameter sets to data are depicted as violin plots. Compared to no intervention (white, right hand side), treating current PWID and decreased injection initiation were associated with the largest predicted reductions to HCV prevalence (A). Reducing injection initiation was associated with the largest predicted reductions to new chronic HCV cases (B).

**Figure 3.**
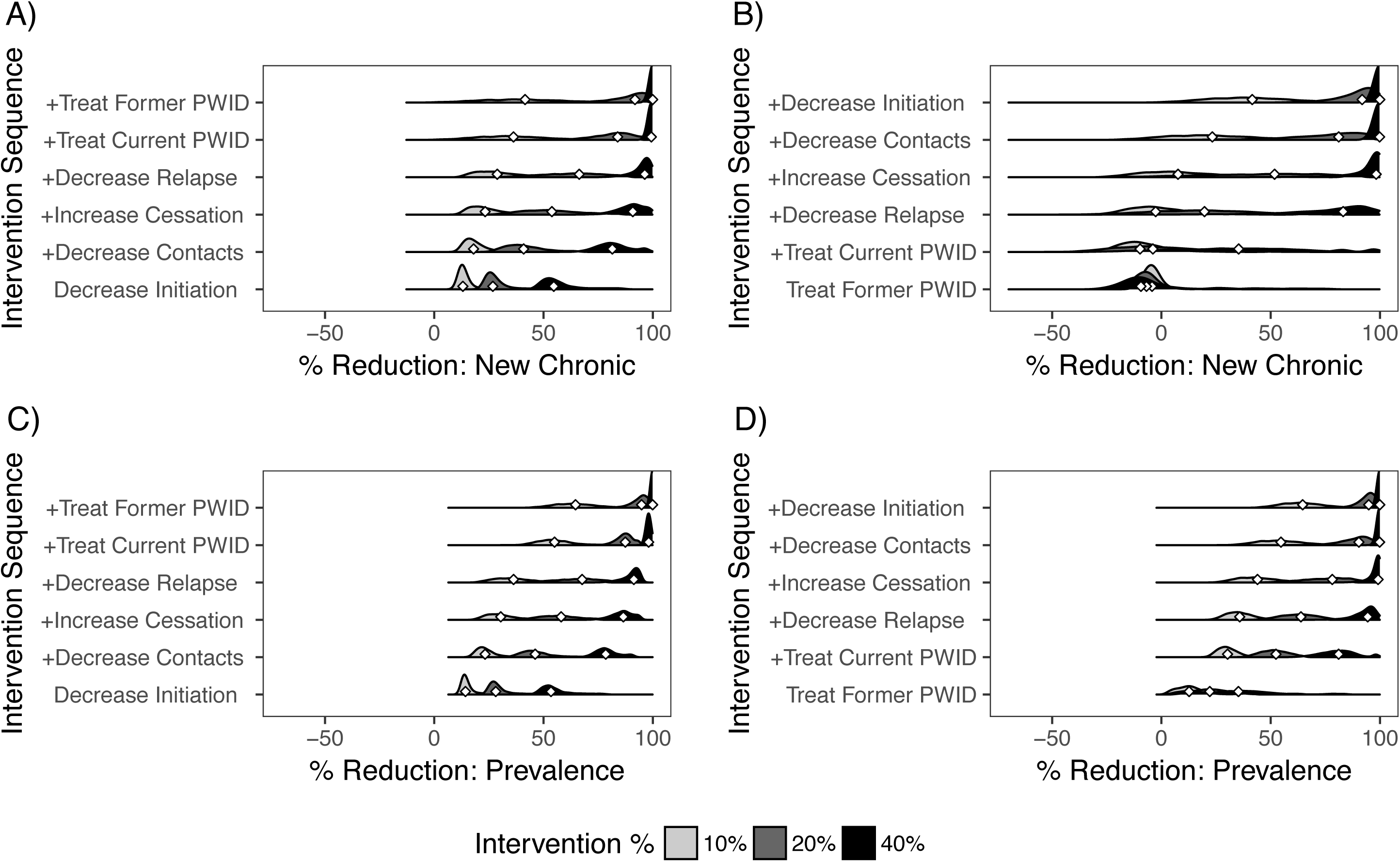
Counterfactual Simulation of Combined Interventions. Histograms of the predicted percent reduction in new chronic cases (panels A and B) and prevalence (panels C and D) after 14-years simulation is plotted for each of 2,500 parameter sets. All percent reduction calculations are calculated in comparison to no intervention and interventions are sequentially added from a base of primary (panels A and C) versus tertiary (panels B and D) interventions. Diamonds denote the median percent reduction. Primary prevention through decreased initiation of non-PWID to injecting was concurrently associated with predicted reductions to new chronic HCV cases (A) and prevalence (B). Reductions continued with the introduction of secondary and tertiary measures. Tertiary measures (HCV treatment) were associated with predicted reductions to prevalence (D) but not new chronic cases (B). Introduction of secondary on top of tertiary prevention measures predicted decreases in new chronic cases, although predicted declines to new chronic cases were smaller with the addition of additional interventions until all primary, secondary, and tertiary interventions were engaged simultaneously. The variability in predicted reductions in new chronic cases (panels A and B) was larger than prevalence reductions (panels C and D). Interventions including treatment with secondary interventions (panels B and D) exhibited higher variability in predicted case counts compared to those with primary and secondary interventions (panels A and C).

Among the best-fitting 2,500 simulations, the largest predicted reduction in chronic HCV prevalence at year 14 of simulation occurred by treating current PWID, although variability in results increased with increasing treatment levels (Figure 2A). Primary prevention interventions (decreased injection initiation) predicted the largest and least variable decreases to new chronic HCV cases across parameter sets, while secondary measures had modest reductions and/or high variability (Figure 2B). New chronic cases were not generally avoided by treating former PWID. Treatment of only former PWID led to small increases or stagnation in predicted new chronic case counts, likely due to reinfection after treatment. Treatment of current or former PWID in the absence of other interventions predicted reductions in new chronic HCV cases in a subset of parameter sets characterized by high injection initiation rates, high numbers of syringe sharing partners, and low rates of relapse after injection cessation. Increased treatment duration and interventions decreasing mortality among current or former PWID did not impact HCV prevalence or new chronic cases (Web Figure 4).

When considered in combination, adding secondary interventions alongside primary (decreased injection initiation) or tertiary prevention (former PWID treatment) predicted similar average reductions in HCV prevalence at year 14 (Figure 3). Strategies including primary prevention measures averted new chronic cases in all scenarios, and predicted reductions increased by adding secondary and tertiary measures. By comparison, predicted reductions in new chronic cases after including secondary measures alongside treatment were smaller. Interventions including treatment with secondary interventions also exhibited higher variability in predicted case counts across parameter sets compared to those with primary and secondary interventions.

## DISCUSSION

We used an HCV transmission model among young PWID informed by Michigan HCV surveillance data to evaluate the relative benefits of primary, secondary and tertiary interventions to reduce HCV prevalence and incidence. Simulation results suggested that primary prevention initiatives to decrease injection initiation among adolescents and young adults would be the most effective HCV intervention strategy. Both singly and in combination with secondary prevention measures, such as decreasing relapse, increasing cessation, and decreasing syringe sharing, primary prevention predicted reductions to both prevalence and new chronic cases. Predicted prevalence reductions were nearly comparable in magnitude to treatment interventions, although treatment of currently injecting PWID was predicted to be most effective in reducing HCV prevalence. In line with the concept of ‘treatment as prevention’, current PWID treatment also yielded moderate reductions to new chronic HCV cases at high levels of treatment, particularly if combined with secondary and primary prevention initiatives. In addition to reducing the public health impacts of HCV directly, multi-strategy interventions including behavioral interventions focused on IDU would concurrently reduce the economic costs of HCV, which increase with earlier age of infection, and the economic and societal costs of heroin and other opioid use disorders (51,52).

The majority of parameter sets used to simulate interventions tended towards the lower ranges of current PWID prevalence, cessation, and syringe sharing contacts among 15-19 year olds and the upper range of lifetime PWID prevalence. Our finding that primary prevention initiatives have the greatest potential for reducing incidence and prevalence together, therefore, are directly generalizable HCV transmission scenarios exhibiting these characteristics. Interestingly, a small number of parameter sets fitting well to data and used to simulate interventions had high injection initiation rates, high numbers of injection partners, and low rates of relapse after injection cessation. Under these conditions, treatment of both current and former PWID was predicted to be highly effective in reducing chronic HCV prevalence and incidence, even in the absence of primary and secondary interventions. Intervention results under each of these two scenarios could be useful in identifying optimal intervention approaches if behavioral characteristics of PWID in a specific locale are known or measured, such as may be done through case or outbreak investigation.

In sensitivity analyses, mortality reduction interventions among current and former PWID did not qualitatively change predicted primary, secondary, and tertiary intervention results, and absolute increases in predicted HCV prevalence and new chronic cases in the presence of mortality reduction interventions were exceedingly small. Additionally, in agreement with Echevarria et al., treatment predictions were insensitive to treatment duration (14). These results are important given that both could theoretically result in maintenance of a larger PWID population, who could be re-infected and/or contribute to HCV transmission. Practically, secondary and tertiary HCV interventions are and should continue to be coupled alongside mortality reduction interventions. For example, harm reduction programs provide co-located services that include provision of sterile injecting equipment, bystander overdose response training, bloodborne virus testing, and referral to treatment for HCV, Human Immunodeficiency Virus, and other services, such as mental health counseling, substance use disorder treatment, or opioid maintenance therapy (53). Additionally, decreased treatment duration makes treatment more feasible, accessible, and less costly.

Like all modeling exercises, we made several simplifying assumptions, and we were limited by existing data. Our model only considers HCV acquisition through IDU among young PWID and results are not generalizable to other age or risk groups. Cases with missing risk factor data were assumed to be PWID, consistent with PWID being the most common risk factor for HCV (3,4,54). Because surveillance data suffers from under-reporting and missing data, we applied a reporting rate developed for acute cases to adjust for under-reporting of new chronic cases because there were very few acute cases reported during the time period under study (47). To address uncertainty in model parameter values, including the reporting rate, we sampled parameters and used values most consistently fitting to surveillance data to simulate interventions.

Our model assumed homogeneous mixing beyond age and used a contact matrix from Mossong et al., a European study of social contacts, that may not reflect the age-related patterns underlying syringe sharing (49). Mossong et al. provided relevant age-group contacts and because we included sampled parameters, we likely capture many plausible syringe sharing scenarios and present summaries of fit to data to show which scenarios were most consistent with observed data (Web Figure 2). Treatment interventions assumed a wholly treatment-naïve population, however this is likely realistic given the historically low treatment rates among PWID during the time period under study (55–58). Further, treatment was simulated as 100% effective and should therefore be interpreted as the maximum possible treatment effect, given that randomized controlled trials support that approximately 89% of patients are cured of HCV by 12 weeks of DAA therapy (59).

## Conclusions

HCV surveillance data is a valuable data source for understanding HCV transmission and identifying local intervention opportunities among young PWID. In Michigan, primary prevention interventions to decrease injection initiation should be a priority to ensure that HCV prevalence does not further increase among young persons. Treatment for current PWID reduces prevalence more than treating former PWID, thus persons at all stages of use or recovery should be connected to HCV treatment alongside primary and secondary prevention interventions.

## ACKNOWLEDGMENTS

Financial support: This work was supported by the Thomas J Francis, Jr. award from the Department of Epidemiology at the University of Michigan School of Public Health and by the Integrated Training in Microbial Systems fellowship at the University of Michigan.

The authors thank Geoff Brousseau, MPH and Kathryn E. Macomber, MPH at the Michigan Department of Health and Human Services for their help in obtaining the surveillance data used in this work.

